# Dietary suppression of MHC-II expression in intestinal stem cells enhances intestinal tumorigenesis

**DOI:** 10.1101/2020.09.05.284174

**Authors:** Semir Beyaz, Jatin Roper, Khristian E. Bauer-Rowe, Michael E. Xifaras, Ilgin Ergin, Lenka Dohnalova, Moshe Biton, Karthik Shekar, Haiwei Mou, Onur Eskiocak, Deniz M. Özata, Katherine Papciak, Charlie Chung, Mohammed Almeqdadi, Miriam Fein, Eider Valle-Encinas, Aysegul Erdemir, Karoline Dogum, Aybuke Garipcan, Hannah Meyer, James G. Fox, Eran Elinav, Alper Kucukural, Pawan Kumar, Jeremy McAleer, Christoph A. Thaiss, Aviv Regev, Stuart H. Orkin, Ömer H. Yilmaz

## Abstract

Little is known about how interactions between diet, immune recognition, and intestinal stem cells (ISCs) impact the early steps of intestinal tumorigenesis. Here, we show that a high fat diet (HFD) reduces the expression of the major histocompatibility complex II (MHC-II) genes in ISCs. This decline in ISC MHC-II expression in a HFD correlates with an altered intestinal microbiome composition and is recapitulated in antibiotic treated and germ-free mice on a control diet. Mechanistically, pattern recognition receptor and IFNg signaling regulate MHC-II expression in ISCs. Although MHC-II expression on ISCs is dispensable for stem cell function in organoid cultures *in vitro*, upon loss of the tumor suppressor gene *Apc* in a HFD, MHC-II- ISCs harbor greater *in vivo* tumor-initiating capacity than their MHC-II+ counterparts, thus implicating a role for epithelial MHC-II in suppressing tumorigenesis. Finally, ISC-specific genetic ablation of MHC-II in engineered *Apc*-mediated intestinal tumor models increases tumor burden in a cell autonomous manner. These findings highlight how a HFD alters the immune recognition properties of ISCs through the regulation of MHC-II expression in a manner that could contribute to intestinal tumorigenesis.

## Introduction

Diet is a major lifestyle factor that influences health and disease states, including cancer (Beyaz and Yilmaz, 2016; Zitvogel et al., 2017). Significant epidemiologic and preclinical studies link obesity to intestinal cancer incidence. However, how the adaptation of the intestinal epithelium to pro-obesity diets alters the cancer risk remains elusive (Basen-Engquist and Chang, 2011; Calle et al., 2003; Gallagher and LeRoith, 2015). The intestinal epithelium is maintained by Lgr5^+^ intestinal stem cells (ISCs) that reside at the crypt base and give rise to the diverse, specialized cell types of the intestinal lining (Barker et al., 2007). These rapidly renewing ISCs coordinate intestinal adaptation in response to environmental inputs such as diet by balancing stem cell self-renewal with differentiation divisions (Beyaz et al., 2016; Wang et al., 2018; Yang et al., 2008; Yilmaz et al., 2012). ISCs are also the cells-of-origin for many early intestinal tumors and lie at the interface of dietary nutrients, commensal microbes and immune cells. Thus, understanding how diet induces changes in ISCs and their surrounding components may shed light on the early steps involved in initiation of intestinal cancers (Barker et al., 2009; Belkaid and Hand, 2014; Clevers, 2013; Hooper et al., 2012; Thaiss et al., 2016).

We recently demonstrated that a pro-obesity high fat diet (HFD) enhances intestinal tumorigenicity in part by boosting intestinal stem cell (ISC) and progenitor cell function and increasing the ability of these cells to undergo oncogenic transformation upon loss of the tumor suppressor gene *Apc* (Beyaz et al., 2016). Because interactions between cancer cells and the immune system influence tumor initiation and progression, it is important to understand the crosstalk between tumor-initiating ISCs and immune cells. Although cancers develop several strategies to evade the immune system (Grasso et al., 2018; Liu et al., 2017), little is known about how diet-induced obesity impacts the interaction of ISCs with the immune system at the start of tumor formation.

## Results

### HFD dampens MHC-II expression in ISCs

To explore how a high fat diet (HFD) perturbs immunomodulatory gene expression in ISCs, we examined our previous mRNA-sequencing (RNA-seq) dataset of Lgr5^+^ ISCs isolated from control and HFD-fed mice (Beyaz et al., 2016). We found that ISCs derived from HFD-fed mice significantly downregulate immunomodulatory genes that are involved in the MHC-II pathway (*H2-Aa, H2-Ab1, Ciita*), anti-microbial response/inflammation (*Reg3g, Nfkb2*) and costimulation (*Icosl, Sectm1a, Sectm1b*) (Choi et al., 2011; Dong et al., 2001; Howie et al., 2013; Mukherjee and Hooper, 2015; Tomas et al., 2016) (Fig. 1A, S1A). Given the recently reported heterogeneity within ISCs(Barriga et al., 2017; Biton et al., 2018b), we next performed single-cell RNA-Seq (scRNA-Seq) of ISCs derived from control or HFD mice. Consistent with the bulk RNA-seq profiles, *H2-Ab1*, a key component of MHC-II complex, was among the top 5 differentially expressed genes with > 3-fold higher expression in control compared to HFD ISCs (Fig. 1B, S1B, C) (see methods, MAST test, p < 1e-10)). To determine the extent of MHC-II downregulation in response to HFD in our scRNA-seq data, we ranked individual ISCs based on their expression pattern of MHC-II pathway genes (MHC-II score, Table 1) and found that HFD ISCs had consistently lower MHC-II score compared to control ISCs (Fig. 1C). We then selected the upper and lower quartiles and performed differential expression analysis between MHC-II low and MHC-II high HFD ISCs. As expected, MHC-II pathway genes (*H2-Ab1, Cd74, H2-Aa, H2-DMa*) were amongst the top upregulated genes in MHC-II high cells. The genes that were upregulated in MHC-II low cells include the fatty acid-binding protein *Fabp2*, chaperones *Dnajc2* and *Hspd1* (Fig. 1D).

**Figure 1.**
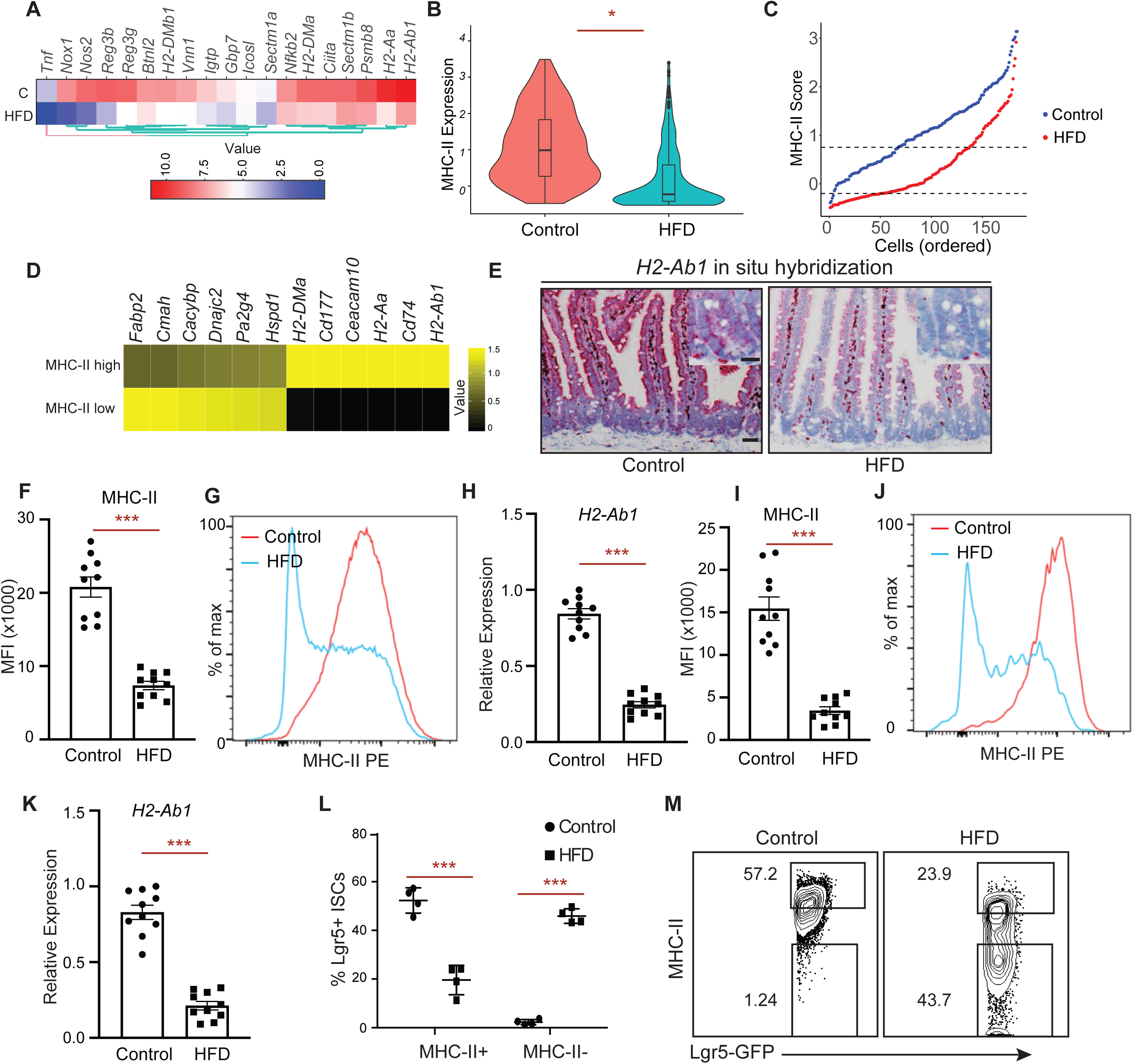
High Fat Diet reduces MHC-II expression on Lgr5-GFP^**hi**^ intestinal stem cells (ISCs). **A**. A heat map of downregulated genes assessed by bulk RNA-seq in Lgr5-GFP^hi^ intestinal stem cells (Lgr5+ ISCs) isolated from long-term high fat diet (HFD)-fed mice compared to control mice (n=2). **B**. Violin plots demonstrating MHC-II expression in single ISCs isolated from control (n=171 cells, 2 independent experiments) or HFD mice by single cell RNA-seq (scRNA-seq) (n=144 cells, 2 independent experiments). **C**. Control (blue) and HFD (red) Lgr5+ ISCs ranked according to their expression of MHC II pathway genes (y-axis). Dashed lines correspond to y-intercepets of −0.2 and 0.75, which are the 25^th^ and 75^th^ percentile of scores in HFD cells, used to define MHC2 low (score < −0.2) and MHC2 high (score > 0.75) HFD cells. In contrast, these values correspond to 1^st^ and 35^th^ percentile of scores in control cells. **D**. Heatmap showing differentially expressed (DE) genes (rows) between MHC2 low and MHC2 high HFD ISCs as defined in panel C. **E**. Single-molecule *in situ* hybridization of MHC-II (*H2-Ab1*) in control and HFD mice (*n*=5). **F, G**. Mean fluorescence intensity (MFI) of MHC-II in Epcam+ cells from crypts of control and HFD mice (**F**, *n*=10 mice). Representative flow cytometry histogram plots of MHC-II expression in Epcam+ cells (**G**). **H**. Relative expression of MHC-II (*H2-Ab1*) in Epcam+ cells isolated from crypts of control and HFD mice (*n*=10 mice). **I, J**. Mean fluorescence intensity (MFI) of MHC-II in Lgr5+ ISCs from crypts of control and HFD mice (**I**, *n*=10 mice). Representative flow cytometry histogram plots of MHC-II expression in Epcam+ cells (**J**). **K**. Relative expression of MHC-II (*H2-Ab1*) in Lgr5+ ISCs isolated from crypts of control and HFD mice (*n*=10 mice). **L, M**. Frequency of MHC-II+ and MHC-II- ISCs in control and HFD mice by flow cytometry (**L**, *n*=4, mean □± □s.d). Representative flow cytometry plots of MHC-II in control and HFD ISCs (**M**). Unless otherwise indicated, data are mean □± □s.e.m. from *n* independent experiments; \**P* □< □0.05, \*\**P* □< □0.01, \*\*\**P* □< □0.001 (Student’s *t*-tests). Scale bars, 50 □μm (**E**) and 20 □μm (**E**, inset).

Although MHC-II expression and function is generally considered to be restricted to professional antigen presenting cells, several studies have demonstrated that intestinal epithelial cells (IECs) express high levels of MHC-II and are able to capture, process and present antigens to CD4+ T cells, including recent work from our group on ISCs (Biton et al., 2018b; Cerf-Bensussan et al., 1984; Dahan et al., 2007; Hershberg et al., 1997; Telega et al., 2000; Thelemann et al., 2014; Westendorf et al., 2009) (Fig. S1F). We validated the reduction of MHC II expression on IECs and ISCs under a HFD in several ways. *In situ* hybridization for *H2-Ab1* showed that MHC-II is expressed in the epithelium of control mice epithelium, but was dampened in HFD mice throughout the epithelium, including in the intestinal crypt where ISCs reside (Fig. 1E). Moreover, by flow cytometry, both Epcam+ IECs (Fig. 1F, G) and Lgr5+ ISCs (Fig. 1I, J) expressed high levels of MHC-II protein on their cell surface at steady state, and this expression was significantly downregulated in response to a HFD. Finally, we confirmed that sorted Epcam+ IECs (Fig. 1H) and Lgr5+ ISCs (Fig. 1K) from HFD-fed mice significantly downregulated MHC-II expression by qPCR. We then partitioned ISCs into two subpopulations based on their MHC-II expression pattern: MHC-II+ and MHC-II- and assessed their frequencies in control or HFD-fed mice. While in control mice most ISCs were MHC-II+, a HFD led to a significant reduction in the frequency of MHC-II+ ISCs and to a concomitant increase in the frequency of MHC-II- ISCs (Fig. 1L, M). We sorted these ISC subpopulations to confirm MHC-II expression levels. Consistent with our scRNA-seq analysis, the expression levels of *H2-Ab1* mRNA was differentially expressed in these subpopulations both in control and HFD-fed mice, with a significant reduction in response to HFD (Fig. S1D). Altogether, these results indicate HFD leads to suppression of MHC-II expression in the intestinal epithelial cells including ISCs.

### PPAR-δ activation or obesity does not affect MHC-II expression in ISCs

A HFD perturbs multiple epithelial-intrinsic and -extrinsic pathways that may influence the regulation of MHC-II expression in ISCs(Fu et al., 2019; Schulz et al., 2014). Because our prior findings demonstrated that a PPAR program mediates many of the effects of a HFD in ISCs (Beyaz et al., 2016; Beyaz and Yilmaz, 2016), we next assessed the activation status of a PPAR-δ program in ISCs based on their MHC-II expression pattern in response to a HFD. We found no difference in the expression levels of signature the PPAR-δ target genes *Hmgcs2* and *Jag1*, or the stem cell marker *Lgr5* in MHC-II+ and MHC-II- ISCs in both diet groups (Fig. S1E). Interestingly, agonist-induced PPAR-d activation also did not reduce MHC-II expression in ISCs, indicating that HFD-induced MHC-II downregulation is independent of PPAR-δ activity in ISCs (Fig. S1I, J). To determine whether MHC-II expression in ISCs is reduced in an independent model of obesity, we examined leptin receptor deficient (db/db) mice, an obesity model that develops on a control diet(Beyaz et al., 2016) (Fig. S1H). Lgr5+ ISCs isolated from both lean control (db/+) and obese (db/db) mice expressed high levels of MHC-II, indicating that diet-induced alterations rather than obesity *per se* inhibits MHC-II expression in ISCs (Fig. S1K, L).

### MHC-II expression in ISCs does not influence organoid-forming capacity

We next assessed whether HFD-mediated downregulation of MHC-II affects stemness. First, we assayed the functional potential of MHC-II+ and MHC-II- ISCs, using an *in vitro* approach based on the ability of isolated ISCs to form organoid bodies in 3-D culture (Beyaz et al., 2016; Sato et al., 2009a). MHC-II expression levels in control and HFD ISCs did not affect *in vitro* organoid formation (Fig. S1F). To further ascertain whether MHC-II expression regulates ISC function, we generated mice with intestinal epithelium-specific MHC-II deletion (vil-iKO; see Methods). Intestinal specific loss of MHC-II did not affect the weight and length of the small intestine or impact crypt depth and villi height (Fig. S2A-G). Furthermore, the proliferation or organoid-forming capacity of ISCs was unaltered upon loss of MHC-II in the intestinal epithelium (Fig. S3A-F). To assess how MHC-II regulates ISC function specifically, we ablated MHC-II in Lgr5^+^ ISCs (Lgr5-iKO; see Methods) and found that loss of MHC-II in ISCs did not affect the capacity of ISCs to initiate organoids (Fig. S3G-H). To determine how MHC-II influences Lgr5+ ISC function in vivo, we performed LacZ lineage tracing analysis and did not detect a significant difference in LacZ+ cells between WT and Lgr5-iKO mice. While these data show that the *in vitro* organoid-forming capacity of ISCs is independent of MHC-II expression and that MHC-II loss does not grossly alter intestinal homeostasis and ISC output (i.e LacZ+ progeny) at steady state, as we previously reported(Biton et al., 2018a) MHC-II may influence ISC differentiation through engaging with T helper cell cytokines in the context of inflammation (Fig. S2I).

### MHC-II expression on ISCs depends on the intestinal microbiome

The intestinal microbiome plays a significant role in regulating intestinal immunity (Belkaid and Hand, 2014; David et al., 2014; Hooper et al., 2012; Round and Mazmanian, 2009; Thaiss et al., 2016). Since dietary perturbations are among the major external factors shaping the intestinal microbiome, we asked whether HFD-induced alterations in the microbiome influence MHC-II expression in ISCs and the intestinal epithelium. Consistent with previous findings, HFD-induced obesity led to microbial dysbiosis with reduced bacterial diversity (Fig. S4A, B) (Ley et al., 2005; Schulz et al., 2014). To determine whether the microbiome is involved in regulation of epithelial MHC-II levels, we treated mice with broad-spectrum antibiotics, which ablated bacterial diversity and massiviely altered community composition (Fig. S4A, B). Notably, antibiotic treatment was accompanied by decreased MHC-II expression in ISCs and the intestinal epithelium (Fig. 2A, B), comparable to that observed in HFD. To further corroborate the role of the intestinal microbiome on MHC-II expression on ISCs, we generated germ-free Lgr5-GFP-IRES-CreERT2 mice. ISCs from germ-free mice exhibited reduced MHC-II expression both at RNA and protein levels compared to specific pathogen-free control mice (Fig. 2C-F, S4C, D).

**Figure 2.**
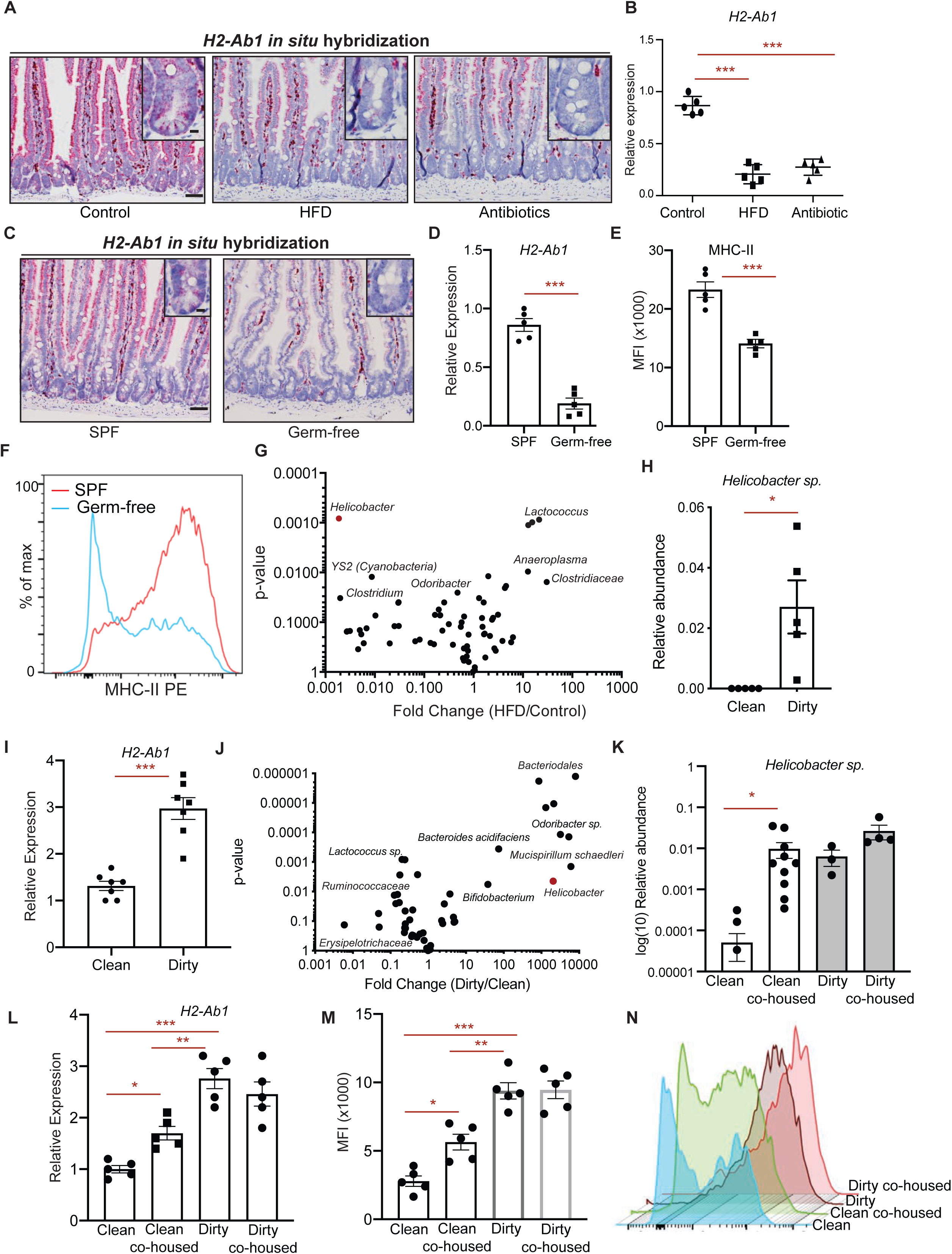
Intestinal microbiome regulates MHC-II expression in ISCs. **A**. *In situ* hybridization for *H2-Ab1* in control, HFD, and antibiotic-treated mice in proximal small intestine (*n=*5). **B**. Relative expression of MHC-II in Lgr5+ ISCs from control, HFD, and antibiotic-treated mice (*n=*5, mean □± □s.d). **C**. Single-molecule *in situ* hybridization of MHC-II (*H2-Ab1*) in specific pathogen-free (SPF) and germ-free mice (*n*=5 mice). **D**. Relative expression of MHC-II (*H2-Ab1*) in Lgr5+ ISCs from SPF and germ-free mice (*n=*5). **E, F**. Mean fluorescence intensity (MFI) of MHC-II in Lgr5+ ISCs from SPF and germ-free mice (**E**, *n*=5). Representative flow cytometry histogram plots of MHC-II expression in Lgr5+ ISCs (**F**). **G**. Volcano plot demonstrating significantly enriched and depleted microbial species in HFD versus control mice (*n*=5). **H**. Relative abundance of *Helicobacter sp*. in mice housed in clean room and dirty room (*n*=5). **I**. Volcano plot demonstrating significantly enriched and depleted microbial species in mice housed in dirty room versus clean room (*n*=6). **J**. Relative expression of MHC-II (*H2-Ab1*) in Epcam+ cells isolated from crypts of mice housed in clean room or dirty room (*n*=7). **K**. Relative abundance of *Helicobacter sp*. in mice housed either in clean room (*n*=10), dirty room (*n*=3), or after co-housing clean mice (*n*=10) with dirty mice (*n*=4) in dirty room. **L**. Relative expression of MHC-II (*H2-Ab1*) in Epcam+ cells isolated from crypts of mice housed either in clean room, dirty room, or after co-housing clean mice with dirty mice in dirty room (*n*=5, ANOVA). **M, N**. Mean fluorescence intensity (MFI) of MHC-II in Epcam+ cells isolated from crypts of mice housed either in clean room, dirty room, or after co-housing clean mice with dirty mice in dirty room (**M**, *n*=5, ANOVA). Representative flow cytometry histogram plots of MHC-II expression in Lgr5+ ISCs (**N**). Unless otherwise indicated, data are mean □± □s.d. from *n* independent experiments; \**P* □< □0.05, \*\**P* □< □0.01, \*\*\**P* □< □0.001 (Student’s *t*-tests).

To gain insight into the spectrum of members of the bacterial microbiome capable of inducing ISC MHC-II expression, we performed comparative 16S rDNA sequencing in HFD-treated mice and controls. Among the bacterial genera most strongly ablated under HFD conditions was *Helicobacter* (Fig. 2G). To determine whether *Helicobacter* colonization in mice correlates with the epithelial MHC-II expression in the intestine, we surveyed our mouse facility for presence or absence of *Helicobacter* species. We determined two separate rooms with mice either naturally colonized with *Helicobacter* species (H+, dirty room) or without detectable *Helicobacter* species (H-, clean room) (Fig. 2H). Notably, MHC-II expression was significantly higher in the intestinal epithelium of mice that are housed in dirty room compared to the mice housed in clean room (Fig. 2I). Comparison of microbial composition between dirty mice and clean mice revealed additional bacteria that are more abundant in dirty mice and associate with high epithelial MHC-II expression (Fig. 2J, S4E). These include *Odoribacter sp*. and *Helicobacter sp*., which were both ablated in response HFD (Fig. 2G, S4E). To determine whether colonization of clean mice with the microbial flora of the dirty mice enhances expression of epithelial MHC-II, we co-housed clean mice with dirty mice in the dirty room (Fig. S4G). While clean and dirty mice have distinct microbial composition at baseline as determined by principal component analysis (PCA), after co-housing the microbial profile of clean mice resembles the dirty mice (Fig. S4G). Importantly, co-housing with dirty mice led to a significant increase in *Helicobacter sp*. abundance in the co-housed clean mice with a concomitant upregulation of epithelial MHC-II compared to mice always remained in the clean room (Fig. 2K-N). These results indicate that MHC-II expression in ISCs is regulated by intestinal commensal bacteria including *Helicobacter sp*., which is altered in response to a HFD.

### Pattern recognition receptor (PRR) and IFN γ signaling regulate MHC-II expression in ISCs

Microbiome-induced activation of pattern recognition receptor (PRR) signaling or proinflammatory cytokines induce MHC-II expression in antigen presenting cells and regulate intestinal homeostasis (Abreu, 2010; Rakoff-Nahoum et al., 2004; van den Elsen, 2011). We surveyed PRR expression patterns in ISCs in control and HFD conditions. Consistent with previous reports (Brown et al., 2014; Caruso et al., 2014; Neal et al., 2012; Price et al., 2018), ISCs expressed several Toll-like receptors (*Tlr1, Tlr2, Tlr3* and *Tlr4*), as well as Nod-like receptors (*Nod1, Nod2*) (Fig. 3A). Among these PRRs, *Tlr2* and *Nod2* were downregulated in response to a HFD (Fig. 3A). To test whether activation of TLR2 and NOD2 pathways is sufficient to increase MHC-II expression in ISCs, we treated mice with a dual TLR2/NOD2 agonist (CL429) (Pavot et al., 2014). Indeed, CL429 treatment led to significant upregulation of MHC-II in both control and HFD ISCs, suggesting that signaling through TLR2 and NOD2 can induce MHC-II expression (Fig. 3B, S5A, B). Of note, CL429 treatment did not fully restore MHC-II levels of HFD-treated mice to control mice, which indicates that additional signaling pathways may be perturbed (Fig. 3B, S5A, B). To determine the necessity of PRR signaling through the adaptor protein Myd88 in regulating MHC-II expression, we utilized intestine-specific Myd88-deficient mice (Myd88 iKO) and found significant downregulation of intestinal MHC-II compared to wild-type controls (Fig. S5C, D).

**Figure 3.**
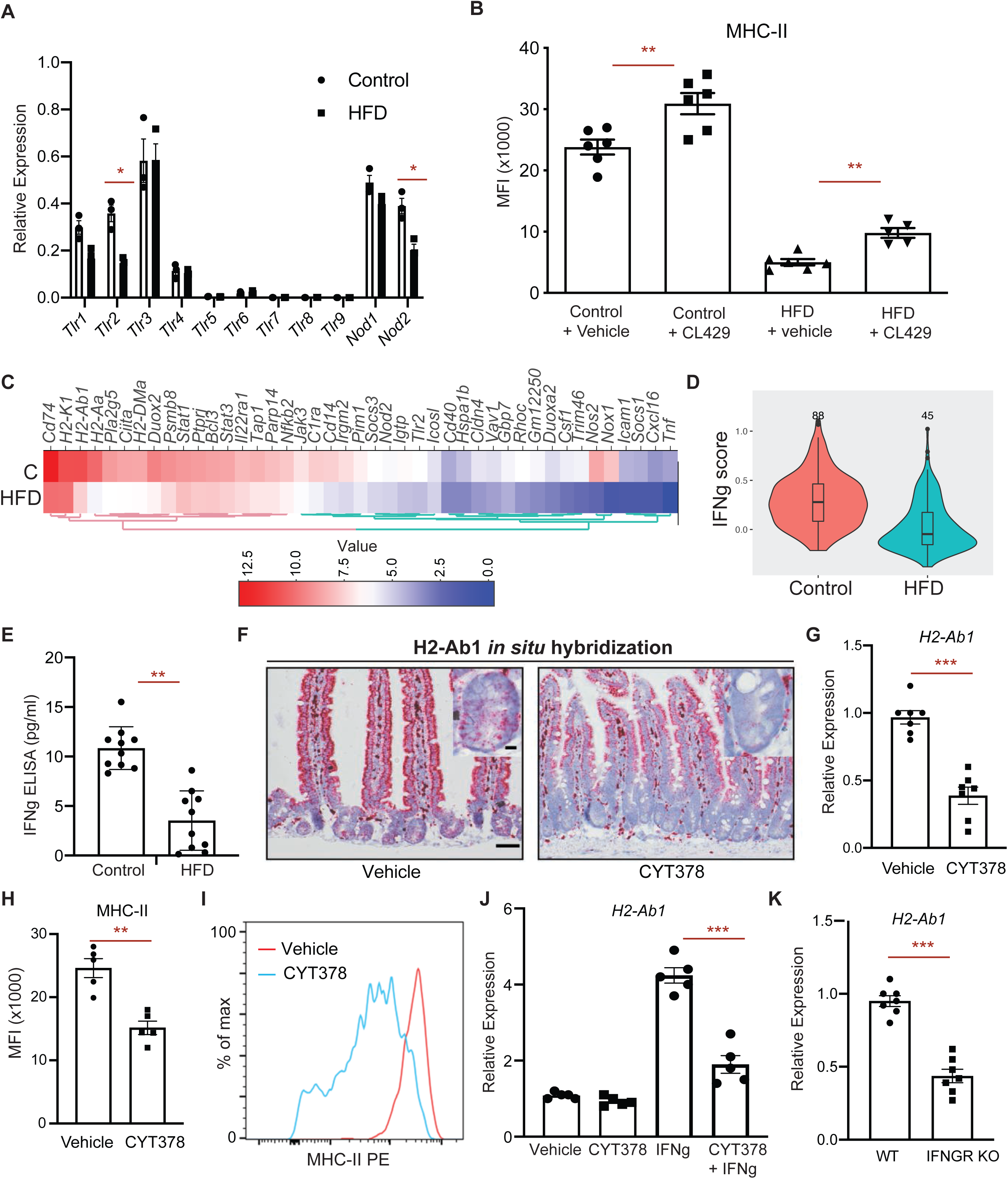
PRR and IFN γ signaling regulate MHC-II expression in ISCs. **A**. Relative expression of pattern recognition receptors (PRR) in control and HFD ISCs (*n*=3). **B**. Mean fluorescence intensity (MFI) of MHC-II in ISCs from vehicle- and TLR2/NOD2 agonist CL429-treated control and HFD mice (*n*=6 mice). **C**. A heat map of expression levels of IFN γ-induced genes between HFD and control ISCs by bulk RNA-seq (*n*=2). **D**. Violin plots demonstrating the expression levels of IFN γ-induced genes in control and HFD ISCs by scRNA-seq. **E**. IFN γ levels in the intestines of control and HFD mice as measured by ELISA (*n*=10). **F**. *In situ* hybridization for *H2-Ab1* in vehicle- and CYT387-treated mice in small intestine (*n=*3). **G**. Relative expression of MHC-II (*H2-Ab1*) in Lgr5+ ISCs from vehicle- and Jak/Stat inhibitor CYT387-treated mice (*n*=7). **H, I**. Mean fluorescence intensity (MFI) of MHC-II in Lgr5+ ISCs from vehicle- and CYT387-treated mice (**H**, *n*=5). Representative flow cytometry histogram plots of MHC-II expression in Lgr5+ ISCs (**I**). **J**. Relative expression of MHC-II (*H2-Ab1*) in intestinal organoids-treated with or without CYT387 and/or IFN γ (*n*=5). **K**. Relative expression of MHC-II (*H2-Ab1*) in Epcam+ cells isolated from crypts of control or IFNGR KO (*n*=5). Unless otherwise indicated, data are mean □± □s.e.m from *n* independent experiments; \**P* □< □0.05, \*\**P* □< □0.01, \*\*\**P* □< □0.001 (Student’s *t*-tests).

The intestinal microbiome also influences the activity of proinflammatory cytokine signaling, such as the proinflammatory cytokine interferon-gamma (IFN γ) and downstream JAK1/2 signaling pathways (van den Elsen, 2011). IFN γ is a potent inducer of MHC-II expression in antigen presenting cells and other cell types including intestinal epithelial cells (Collins et al., 1984; Thelemann et al., 2014; Wong et al., 1984). We examined the responsiveness to IFNγ of MHC-II+ and MHC-II- HFD ISCs, which express similar levels of the IFNγ receptors *Ifngr1* and *Ifngr2* (Fig S1E). IFNγ treatment led to upregulation of MHC-II in both MHC-II+ and MHC-II- ISCs (Fig. S5E), suggesting that although HFD ISCs express lower levels of MHC-II, they are still responsive to IFNγ stimulation (Choi et al., 2011; Thelemann et al., 2014). To determine whether HFD leads to suppression of epithelial MHC-II expression through dampening IFN γ signaling, we assessed the expression levels of IFN γ-induced genes in control and HFD ISCs. We found ISCs significantly downregulate IFN γ-induced genes, including MHC-II pathway genes (*H2-Ab1, H2-Aa, Ciita, Cd74, H2-DMa*) and upstream pathway genes that regulate MHC-II expression (*Stat1, Stat3, Nfkb2, Jak3*) in response to HFD (Fig. 3C, D, S5F, G). Of note, the levels of IFNγ were significantly reduced in the intestines of HFD mice (Fig. 3E). Notably, administration of a potent JAK1/2 & TBK1/IKKε inhibitor (CYT378) that inhibits both STAT and NFκB signaling to mice significantly downregulated MHC-II expression in ISCs compared to vehicle-treated controls (Fig. 3E-H, S5H, I) (Tyner et al., 2010). We next asked whether IFN γ-induced epithelial MHC-II expression is inhibited by CYT378 in *in vitro* organoid assays. Strikingly, CYT378 blunted the induction of MHC-II expression in response to IFN γ (Fig. 3J). We assessed the necessity of IFN γ-signaling in regulating MHC-II expression by utilizing IFNGR knockout mice and found dampened epithelial MHC-II expression in IFNGR knockout mice compared to wild-type controls (Figure 3K). HFD-mediated reduction in IFN γ-inducible and MHC-II pathway gene expression in ISCs posits the possibility that intestinal immune cells may be depleted in the intestinal epithelium. Indeed, we found significant reduction in CD45+ immune cells infiltration to the crypt epithelium, including CD3+, CD8+ and CD4+ T cells (Figure S6A-H). Altogether, these results suggest that PRR and IFN γ signaling drive MHC-II expression in the intestinal epithelial cells including ISCs and that HFD attenuates MHC-II expression in ISCs by perturbing these pathways. Our data is consistent with previous studies that demonstrated reduced inflammation and immune infiltration in the intestinal crypt in response to a HFD (Beyaz et al., 2016; Johnson et al., 2015; Schulz et al., 2014).

### Dampening MHC-II expression in premalignant ISCs increases intestinal tumorigenesis

Recognition of antigens by T cells through antigen presentation pathways is a major mechanism for triggering anti-tumor immunity (Vanneman and Dranoff, 2012). Dampening the expression of genes involved in antigen presentation to evade anti-tumor immune responses is a hallmark of many cancers and correlates with poor prognosis (Lovig et al., 2002; Rimsza et al., 2004; Tarafdar et al., 2017). We found that adenomas arising in HFD-fed mice upon loss of *Apc* expressed less MHC-II than controls (Fig. S7A, B). To test whether MHC-II expression patterns influence the tumor initiation potential of premalignant ISCs, we studied Lgr5-CreERt2;Apc^L/L^ mice fed a control or HFD diet prior to tamoxifen-induced inactivation of the tumor suppressor gene *Apc*. Whereas premalignant APC^null^ ISCs in control mice were mostly MHC-II+, APC^null^ MHC-II- ISCs in HFD mice expressed significantly less MHC-II both at RNA and protein levels (Fig. S7C-F). Interestingly, as we observed in the non-neoplastic intestine (Fig. S3G, H), MHC-II expression levels on APC^null^ ISCs did not affect their ability to form adenomatous organoids *in vitro* (Fig. 4A, B). This indicates that decreased MHC-II expression has no effect on the cell-intrinsic oncogenic potential of premalignant APC^null^ ISCs in organoid cultures *in vitro*.

**Figure 4.**
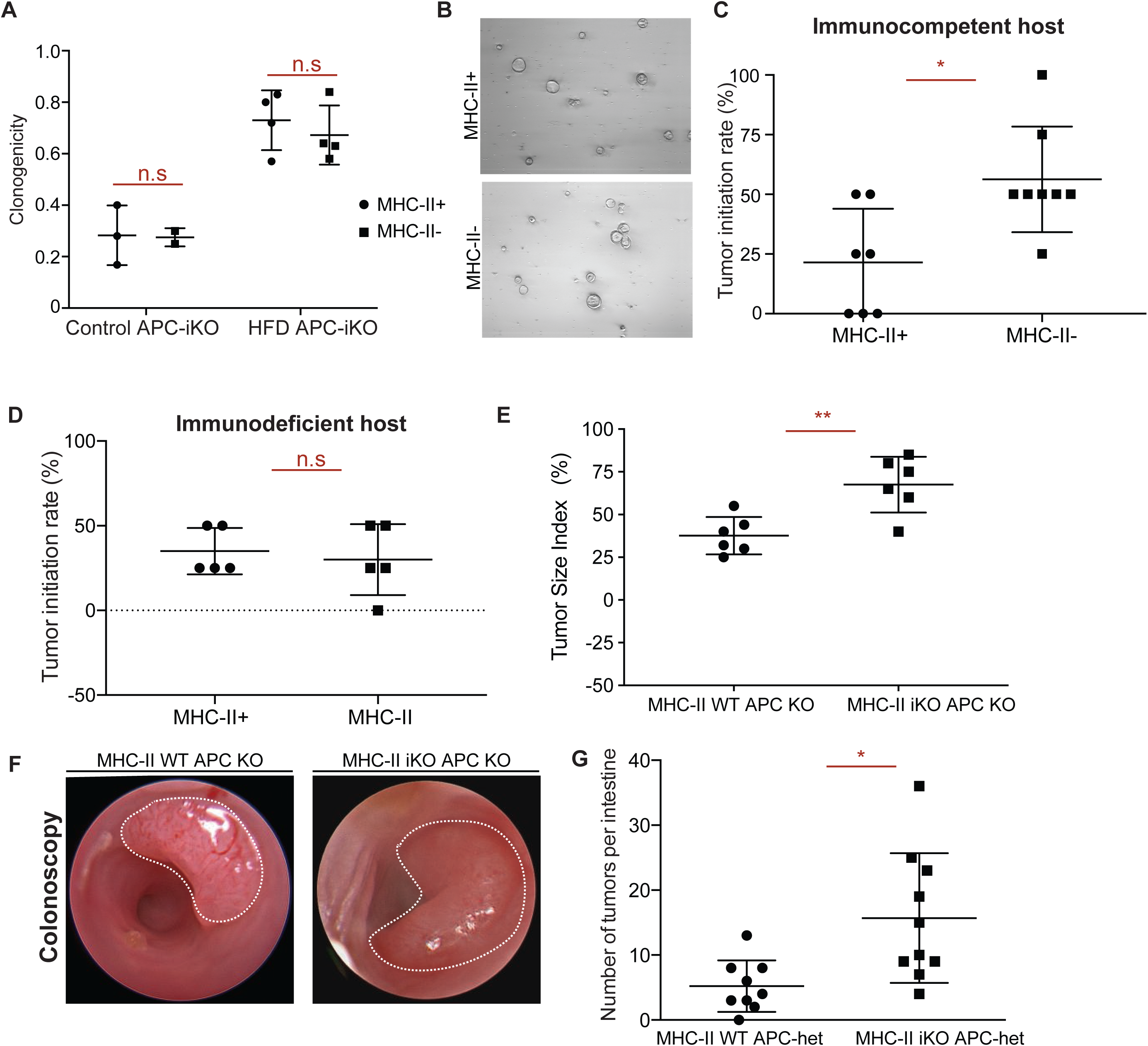
Loss of MHC-II in premalignant ISCs increases tumor initiation. **A, B**. Organoid-initiating capacity of control and HFD MHC-II+ and MHC-II- *Apc*-null ISCs at day 5 (control MHC-II+, *n*=3, control MHC-II-, *n*=3, HFD MHC-II+, *n*=4, HFD MHC-II-, *n=*4). Representative images of HFD MHC-II+ and MHC-II- *Apc*-null organoids at day 5 (**B**). **C**. Tumor initiation rate of orthotopically transplanted MHC-II+ and MHC-II- *Apc*-null ISCs from HFD mice into immunocompetent syngeneic hosts (*n=*8). **D**. Tumor initiation rate of orthotopically transplanted MHC-II+ and MHC-II- *Apc*-null ISCs from HFD mice into immunodeficient (Rag2 KO) hosts (*n=*8). **E, F**. Tumor size index in distal colon of mice that received tamoxifen through endoscopy guided tamoxifen injection to induce tumor formation upon loss of APC (**E**, *n*=6, MHC-II WT APC KO: villin-Cre-ERt2 APC L/L, MHC-II L/+, MHC-II iKO APC KO: villin-Cre-ERt2 APC L/L, MHC-II L/L). Representative optical colonoscopy images of tumors (**F**). **G**. Number of tumors per small intestine in Lgr5-CreERt2 APC^+/-^ MHC-II^+/-^ (*n*=9, MHC-II WT APC-het) and Lgr5-GFP+ APC^+/-^ MHC^-/-^ (*n*=9, MHC-II iKO APC KO) mice 5 months post tamoxifen injection. Unless otherwise indicated, data are mean □± □s.e.m from *n* independent experiments; \**P* □< □0.05, \*\**P* □< □0.01, \*\*\**P* □< □0.001 (Student’s *t*-tests).

To test whether HFD-mediated downregulation of MHC-II in ISCs has pro-tumorigenic effects *in vivo* under immune surveillance, we utilized a recently developed orthotopic, syngeneic colon transplantation assay in mice (Beyaz et al., 2016). We sorted APC^null^ MHC-II+ or APC^null^ MHC-II- Lgr5-GFP^hi^ cells by flow cytometry and transplanted them into the colonic submucosa of syngeneic, immune-competent mice. In contrast to the *in vitro* organoid assay, APC^null^ MHC-II- Lgr5-GFP^hi^ cells exhibited greater tumorigenicity when transplanted *in vivo* than their APC^null^ MHC-II+ counterparts (Fig. 4C, S7G-I). Notably, when transplanted to immunodeficient hosts that lack adaptive immune cells, both MHC-II+ and MHC-II- APC^null^ premalignant cells gave rise to equal numbers of tumors, highlighting the significance of MHC-II recognition by immune cells in controlling intestinal tumor initiation (Fig. 4D, S7J-L). Finally, to ascertain the role of epithelial MHC-II expression during intestinal tumorigenesis, we generated two genetic mouse models. First, using epithelial-specific inducible deletion model of MHC-II together with tumor suppressor *Apc* (vilCreERt2; Apc^L/L^; MHC-II^L/L^), we initated single tumors in the distal colon by administering 4-OHT with our endoscopy-guided injection system. We found that mice with MHC-II KO allele (vil-Cre-ERt2 APC^L/L^, MHC-II^L/+^) gave rise to larger tumors compared to mice with WT allele (vil-Cre-ERt2 APC^L/L^, MHC-II^L/+^) (Fig 4E, F). Second, we initiated tumors using Lgr5+ ISC-specific inducible deletion model of MHC-II together with one copy of the tumor suppressor *Apc* that leads to intestinal tumor formation due to loss of heterozygosity. While we did not detect significant difference in the numbers of T cells inflitrating the tumors at the time of analysis, we found that specific loss of MHC-II in ISCs was associated with greater numbers of intestinal tumors compared to their MHC-II- proficient counterparts (Fig. 4G, S7M,O). These results illustrate that HFD-mediated reduction of MHC-II in premalignant ISCs enhances tumor initiation.

## Discussion

ISCs self-renew and differentiate into cells that constitute the intestinal epithelium, which is continuously replenished (Clevers, 2013). Acquisition of oncogenic mutations in these rapidly cycling stem cells leads to tumors that are subject to clearance by T cells (Agudo et al., 2018; Barker et al., 2009; Schepers et al., 2012). It is also becoming increasingly evident that the crosstalk between tissue stem cells and immune cells influences differentiation, homeostasis and cancer risk (Ali et al., 2017; Biton et al., 2018b; Chakrabarti et al., 2018; Hoytema van Konijnenburg et al., 2017; Lindemans et al., 2015; Naik et al., 2018; Naik et al., 2017). Here, we find that a HFD dampens MHC-II expression in ISCs which can promote intestinal cancer by impacting immune surveillance through MHC II. While MHC-I antigen presentation pathway-mediated activation of cytotoxic CD8^+^ T cells is frequently studied in the context of anti-tumor immune responses, MHC-II- mediated activation of CD4^+^ T cells is also pivotal for tumor immunity (Hung et al., 1998; Takeuchi and Saito, 2017; Tran et al., 2014; Wang, 2001; Xie et al., 2010; Zhang et al., 2009). Indeed, recent studies have demonstrated that MHC-II- restricted CD4^+^ T cells are able to eradicate tumors both directly and indirectly through the licensing of dendritic cells and helping CD8^+^ T cell responses (Haabeth et al., 2016; Hirschhorn-Cymerman et al., 2012; Kreiter et al., 2015; Lu et al., 2017; Spitzer et al., 2017; Tran et al., 2014). Consistent with previous studies demonstrating that tumors downregulate MHC-II expression to escape from immune surveillance (Park et al., 2017; Tarafdar et al., 2017), our data implicate dietary regulation of MHC-II expression in ISCs as playing a critical role in intestinal tumorigenesis. Whether diet-induced reduction in MHC-II expression in ISCs directly or indirectly affects anti-tumor CD4^+^ T cell responses warrants further investigation.

We previously demonstrated that the activation of PPAR-d by the lipid constituents of a HFD is an ISC cell-intrinsic mechanism that enhances tumorigenicity in the intestine (Beyaz et al., 2016). The findings presented here indicate that downregulation of MHC-II in ISCs in response to a HFD provides an orthogonal mechanism contributing to intestinal cancer. Indeed, a recent large scale genome-wide variant scan for colorectal cancer identified variants associated with cancer risk in MHC-II gene loci(Huyghe et al., 2018). Together, our results highlight how diet impacts multiple mechanisms that modify cancer risk in the intestine. Because a HFD influences cancer incidence in both mucosal and non-mucosal tissues (Beyaz et al., 2016; Chen et al., 2018; Kroenke et al., 2013; Pascual et al., 2017; Schulz et al., 2014; Yang et al., 2017; Yang et al., 2008), it will be important to explore whether MHC-II mediated immune surveillance of stem cells also takes place in other tissues and whether it is affected by dietary perturbations.

## Materials and Methods

### Mice, High Fat Diet, and drug treatment

Mice were housed in the Cold Spring Harbor Laboratory and Koch Institute for Integrative Cancer Research. The following strains were obtained from the Jackson Laboratory: *Lgr5-EGFP-IRES-CreERT2* (strain name: B6.129P2-Lgr5^tm1(cre/ERT2)Cle^/J, stock number 008875), *Rosa26-lacZ* (strain name: B6.129S4-Gt(ROSA)26Sor^tm1Sor^/J, stock number 003474), *db/db* (strain name: B6.BKS(D)-Lepr^jb^/J, stock number 000697), *Mhc*^*L/L*^ (strain name: B6.129×1-H2-Ab1^tm1Koni^/J, stock number 013181), IFNGR KO (C57BL/6N-Ifngr1tm1.2Rds/J, stock number 025545. *Apc*^*loxp exon 14*^ (Apc^L/L^) has been previously described(Colnot et al., 2004). *Villin-CreERT2* was a gift from Sylvie Robine. Diet-induced obesity studies were performed by using a high fat diet consisting of 60 kcal% fat (Research Diets D12492) beginning at the age of 8-12 weeks and extending for 9 to 14 months. Control mice were age- and sex-matched and were fed matched purified control diet (Research Diets, D12450J). GW501516 (Enzo) was reconstituted in DMSO at 4.5□mg□ml^−1^ and diluted 1:10 in a solution of 5% PEG400 (Hampton Research), 5% Tween80 (Sigma), 90% H_2_O (injection buffer) for a daily intraperitoneal injection of 4mg kg^−1^. *Apc* exon 14 was excised by tamoxifen suspended in sunflower seed oil (Spectrum S1929) at a concentration of 10 mg ml^-1^ and 250 μl per 25g of body weight, and administered by intraperitoneal injection twice over 4 days before harvesting tissue. BrdU (Sigma) was prepared at 10 mg ml^-1^ in PBS, passed through 0.22μm filter and injected at 100mg kg^-1^. CYT387 (SelleckChem) was reconstituted in DMSO at 10mg ml^-1^ and diluted 1:100 in injection buffer for a daily gavage of 25mg kg^−1^ for 7 days. CL429 (InvivoGen) was reconstituted in DMSO at 5mg ml^-1^ and diluted 1:20 in a solution of injection buffer for a daily gavage of 2mg kg^-1^ for 7 days. For broad-spectrum antibiotic treatment, mice received a mixture of vacomycin (0.5g/l), ampicillin (1g/l), metronidazole (1g/l) and neomycin (1g/l) in the drinking water.

### Immunohistochemistry (IHC) and immunofluorescence (IF)

As previously described(Yilmaz et al., 2012), tissues were fixed in 10% formalin, paraffin embedded and sectioned. Antigen retrieval was performed with Borg Decloaker RTU solution (Biocare Medical) in a pressurized Decloaking Chamber (Biocare Medical) for 3 minutes. Antibodies used: rat anti-BrdU (1:2000, Abcam 6326), mouse monoclonal β-catenin (1:100, BD Biosciences 610154), rabbit monoclonal OLFM4 (1:10,000, gift from CST, clone PP7), Biotin-conjugated secondary donkey anti-rabbit or anti-rat antibodies were used from Jackson ImmunoResearch. The Vectastain Elite ABC immunoperoxidase detection kit (Vector Labs) followed by Dako Liquid DAB+ Substrate (Dako) was used for visualization. All antibody incubations involving tissue or sorted cells were performed with Common Antibody Diluent (Cell Signaling).

### *In situ* hybridization

Single-molecule *in situ* hybridization was performed to detect MHC-II (H2-Ab1, #414731) using Advanced Cell Diagnostics RNAscope 2.5 HD Detection Kit following manufacturer’s instructions.

### Flow cytometry and isolation of ISCs and Paneth cells

As previously reported and briefly summarized here, small intestines and colons were removed, washed with cold PBS-/-, opened laterally and cut into 3-5mm fragments. Pieces were washed multiple times with cold PBS-/-until clean, washed 2-3 with PBS-/-/EDTA (10mM), and incubated on ice for 90-120 minutes while mixing at 30-minute intervals. Crypts were then mechanically separated from the connective tissue by shaking, and filtered through a 70-µm mesh into a 50-ml conical tube to remove villus material (for small intestine) and tissue fragments. Crypts were removed from this step for crypt culture experiments and embedded in Matrigel™ with crypt culture media. For ISC isolation, the crypt suspensions were dissociated to individual cells with TrypLE Express (Invitrogen). Cell labeling consisted of an antibody cocktail comprising IA-PE (eBioscience, 30-F11, 1:500), EPCAM APC (eBioscience, G8.8, 1:100), CD11b-APC-Cy7 (eBioscience, 1:500), and CD45 Pacific Blue (eBioscience, 1:500). Dead cells were excluded from the analysis with the viability dye 7-AAD (Life Technologies). ISCs were isolated as Lgr5-EGFP^hi^Epcam^+^ CD45^−^7-AAD^−^ with a BD FACS Aria II SORP cell sorter into supplemented crypt culture medium for culture.

### Culture media for crypts and isolated cells

Isolated crypts were counted and embedded in Matrigel™ (Corning 356231 growth factor reduced) at 5–10 crypts per μl and cultured in a modified form of medium as described previously(Sato et al., 2009b). Unless otherwise noted, Advanced DMEM (Gibco) was supplemented by EGF 40□ng□ml^−1^ (R&D), Noggin 200□ng□ml^−1^ (Peprotech), R-spondin 500□ng□ml^−1^ (R&D or Sino Biological), *N*-acetyl-L-cysteine 1□μM (Sigma-Aldrich), N2 1X (Life Technologies), B27 1X (Life Technologies), Chiron 10 μM (Stemgent), Y-27632 dihydrochloride monohydrate 20□ng ml^−1^ (Sigma-Aldrich). 25□μL drops of Matrigel™ with crypts were plated onto a flat bottom 48-well plate (Corning 3548) and allowed to solidify for 20□minutes in a 37°C incubator. Three hundred microliters of crypt culture medium was then overlaid onto the Matrigel™, changed every three days, and maintained at 37°C in fully humidified chambers containing 5% CO_2_. Clonogenicity (colony-forming efficiency) was calculated by plating 50–300 crypts per well and assessing organoid formation 3–7□days or as specified after initiation of cultures.

Isolated ISCs or progenitor cells were centrifuged for 5□minutes at 250g, re-suspended in the appropriate volume of crypt culture medium (500–1,000□cells□μl^−1^), then seeded onto 25-30 μl Matrigel™ (Corning 356231 growth factor reduced) containing 1□μM Jagged (Ana-Spec) in a flat bottom 48-well plate (Corning 3548). Alternatively, ISCs and Paneth cells were mixed after sorting in a 1:1 ratio, centrifuged, and then seeded onto Matrigel™. The Matrigel™ and cells were allowed to solidify before adding 300 μl of crypt culture medium. The crypt media was changed every second or third day. Organoid bodies were quantified on days 3, 7 and 10 of culture, unless otherwise specified. In secondary experiments, individual primary organoids were mechanically dissociated and replated, or organoids were dissociated for 10 minutes in TrypLE Express at 32°C, resuspended with SMEM (Life Technologies), centrifuged and resuspended in cold SMEM with viability dye 7-AAD. Live cells were sorted and seeded onto Matrigel™ as previously described.

### Bulk RNA-Seq

#### RNA Isolation

For RNA-seq, total RNA was extracted from 200K sorted Lgr5-GFP^hi^ ISCs or Lgr5-GFP^low^ progenitors by pooling two to five 71-week old male mice per HFD and HFD-control using Tri Reagent (Life Technologies) according to the manufacturer’s instructions, except for a overnight isopropanol precipitation at –20°C. From the total RNA, poly(A)^+^ RNA was selected using Oligo(dT)_25_-Dynabeads (Life technologies) according to the manufacturer’s protocol.

#### RNA-Seq Library Preparation

Strand-specific RNA-seq libraries were prepared using the dUTP-based, Illumina-compatible NEXTflex Directional RNA-Seq Kit (Bioo Scientific) according to the manufacturer’s directions. All libraries were sequenced with an Illumina HiSeq 2000 sequencer.

#### Processing of RNA-seq reads and measuring expression level

Raw stranded reads (40 nt) were trimmed to remove adapter and bases with quality scores below 20, and reads shorter than 35 nt were excluded. High-quality reads were mapped to the mouse genome (mm10) with TopHat version 1.4.1 (Trapnell et al., 2009), using known splice junctions from Ensembl Release 70 and allowing at most 2 mismatches. Genes were quantified with htseq-count (with the “intersect strict” mode) using Ensembl Release 70 gene models. Gene counts were normalized across all samples using estimateSizeFactors() from the DESeq R/Bioconductor package(Anders and Huber, 2010). Differential expression analysis was also performed between two samples of interest with DESeq.

### qRT-PCR

Cells were sorted into Tri Reagent (Life Technologies) and total RNA was isolated according to the manufacturer’s instructions with following modification: the aqueous phase containing total RNA was purified using RNeasy plus kit (Qiagen). RNA was converted to cDNA with cDNA synthesis kit (Bio-Rad). qRT-PCR was performed with SYBR green master mix (Bio-Rad) on Bio-Rad iCycler RT-PCR detection system. For low cell numbers (<1000), qRT-PCR was performed after sequence specific pre-amplification as described in single-cell gene expression analysis. Primers used are listed on Supplementary Table 1.

### Single cell RNA-Seq

#### Cell sorting

FACS (Astrios) was used to sort one single cell into each well of a 96-well PCR plate containing 5µl of TCL buffer with 1% 2-mercaptoethanol. The cells were stained for 7AAD^-^ (Life Technologies), CD45^-^ (eBioscience), CD31^-^ (eBioscience), Ter119^-^ (eBioscience), EpCAM^+^ (eBioscience). To enrich for specific IEC populations, cells were isolated from control or HFD Lgr5-GFP mice, stained with the antibodies mentioned above and gated for GFP-high (stem cells) and GFP-low (TAs). A population control of 200 cells was sorted into one well and a no-cell control was sorted into another well. After sorting, the plate was sealed tightly with a Microseal F and centrifuged at 800g for 1 min. The plate was immediately frozen on dry ice and kept at −80°C until ready for the lysate cleanup.

#### Plate-based scRNA-seq

Libraries were prepared using a modified SMART-Seq2 protocol as previously reported (Picelli et al., 2014). Briefly, RNA lysate cleanup was performed using RNAClean XP beads (Agencourt), followed by reverse transcription with Maxima Reverse Transcriptase (Life Technologies) and whole transcription amplification (WTA) with KAPA HotStart HIFI 2× ReadyMix (Kapa Biosystems) for 21 cycles. WTA products were purified with Ampure XP beads (Beckman Coulter), quantified with Qubit dsDNA HS Assay Kit (ThermoFisher), and assessed with a high sensitivity DNA chip (Agilent). RNA-seq libraries were constructed from purified WTA products using Nextera XT DNA Library Preparation Kit (Illumina). On each plate, the population and no-cell controls were processed using the same method as the single cells. The libraries were sequenced on an Illumina NextSeq 500.

#### Computational analysis of scRNA-seq

We profiled Lgr5-high ISCs sorted from control (n=192 cells) and HFD (n=192 cells) using a full length scRNA-seq method (Picelli et al., 2014). Each condition included two replicate 96-well plates from 2 different mice. Expression levels of gene loci were quantified using RNA-seq by Expectation Maximization (RSEM) (Li and Dewey, 2011). Raw reads were mapped to a mouse transcriptome index (mm10 UCSC build) using Bowtie 2 (Langmead and Salzberg, 2012), as required by RSEM in its default mode. On average, 90% of the reads mapped to the genome in every sample. and 55% of the reads mapped to the transcriptome. RSEM yielded an expression matrix (genes x samples) of inferred gene counts, which was converted to TPM (transcripts per million) values and then log-transformed after the addition of 1 to avoid zeros. After filtering cells with low QC metrics (< 400,000 mapped reads, transcriptomic mapping rate < 35% and < 1500 genes detected), we selected 171 control cells and 144 HFD cells for further analysis.

We identified 379 highly variable genes using Seurat’s MeanVarPlot function. H2-Ab1, a key component of MHC-II complex, was among the top 5 differentially expressed genes with > 3-fold higher expression in control compared to HFD ISCs as assessed using MAST test.(Finak et al., 2015) A Gene Ontology analysis of these genes against a background of genes matched in average expression levels showed an enrichment of terms consistent with intestinal biology such as arachidonic acid metabolic process (GO:0019369), intestinal absorption (GO:0050892), as well as immune response, such as “antigen processing and presentation of exogenous peptide antigen” (GO:0002478) and “defense response to Gram-negative (GO:0050829), and Gram-positive (GO:0050830) bacterium”. We performed principal component analysis (PCA) on the data based on the variable genes, and embedded 10 statistically significant PCs identified using a permutation test (Shekhar et al., 2016) on a 2D map using t-distributed stochastic neighbor embedding (tSNE).

### DQ-Ovalbumin Assay

Lgr5-GFP^hi^ intestinal stem cells were plated in Matrigel and crypt media for one day. The following day, the crypt media was replaced with crypt media containing 20 µg/mL DQ-ovalbumin (Invitrogen) and cells were incubated at 4C and 37C for 24 hours. Cells were harvested by washing three times in PBS, removing the Matrigel using Cell Recovery Solution (Corning) and filtered through a 40 µm mesh. Mean fluorescent intensity was analyzed using CellSimple (Cell Signaling).

### Orthotopic transplantation

*Apc*^*L/L*^; *Lgr5-EGFP-IRES-CreERT2* mice were injected with two doses of tamoxifen I.P. Four days later, *Apc*-null Lgr5-GFP^hi^ ISCs and Lgr5-GFP^low^ progenitors were sorted by flow cytometry, as described above. For primary cell transplantations, 10,000 *Apc*-null Lgr5-GFP^hi^, MHC^hi^ and MHC^low^ ISCs were resuspended into 90% crypt culture media (as described) and 10% Matrigel™, and then transplanted into the colonic lamina propria of C57BL/6 recipient mice as previously described (Beyaz et al., 2016; Roper et al., 2017). Mice then underwent colonoscopy eight weeks later to assess tumor formation. Colonoscopy videos and images were saved for offline analysis. Following sacrifice, the distal colons were excised and fixed in 10% formalin, then examined by hematoxylin and eosin section to identify adenomas. Histology images were reviewed by gastrointestinal pathologists who were blinded to treatment groups.

### Taxonomic Microbiota Analysis

Frozen fecal samples were processed for DNA isolation using the Qiagen PowerSoil kit according to the manufacturer’s instructions. 1 ng of purified fecal DNA was used for PCR amplification. Amplicons spanning the variable region 1/2 (V1/2) of the 16S rRNA gene were generated by using the following barcoded primers: Fwd 5’-XXXXXXXXAGAGTTTGATCCTGGCTCAG-3’, Rev 5’-TGCTGCCTCCCGTAGGAGT-3’, where X represents a barcode base. The reactions were subsequently pooled and cleaned, and the PCR products were then sequenced on an Illumina MiSeq with 500 bp paired-end reads. The reads were then processed using the QIIME (Quantitative Insights Into Microbial Ecology, http://www.qiime.org) analysis pipeline. Rarefaction was used to exclude samples with insufficient count of reads per sample. Sequences sharing 97% nucleotide sequence identity in the 16S region were binned into operational taxonomic units (97% ID OTUs). For beta-diversity, unweighted UniFrac measurements were plotted according to the two principal coordinates based on >10,000 reads per sample.

---

**Supplementary Figure 1.**
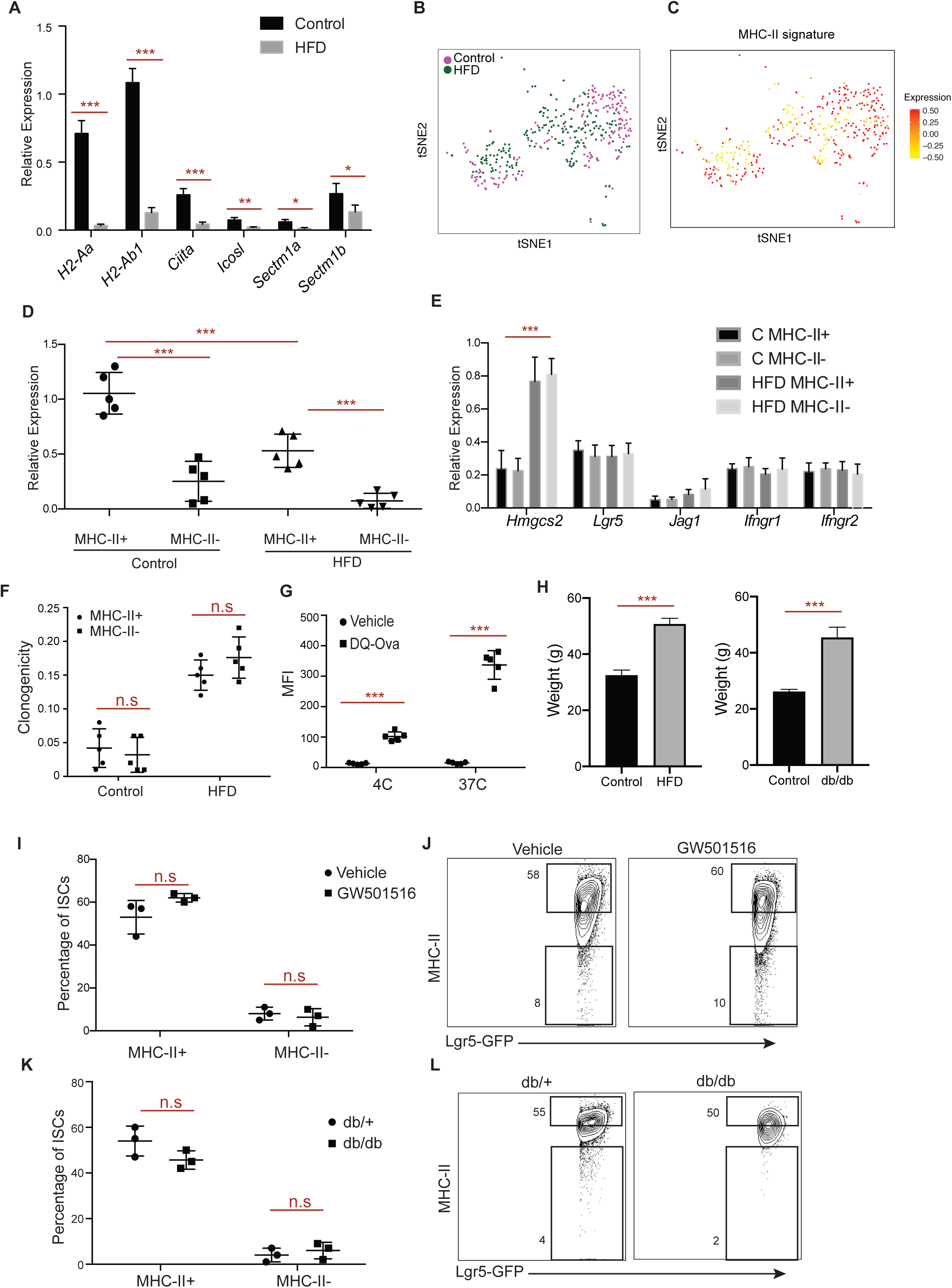
Characterization of the MHC-II+ and MHC-II- ISCs. **A**. Relative expression of immunomodulatory genes in control and HFD Lgr5-GFP^hi^ ISCs (*n*=5). **B**. *t*-Distributed stochastic neighbour embedding (tSNE) analysis of single ISCs isolated from control (n=171 cells, 2 independent experiments) or HFD mice (n=144 cells, 2 independent experiments). **C**. tSNE analysis of single cells using MHC-II pathway signature genes **D**. Relative expression of MHC-II in control and HFD MHC-II+ and MHC-II- ISCs (*n*=5). **E**. Relative expression of signature PPAR-d target (*Hmgcs2*), stem cell marker (*Lgr5*), PPAR-d-dependent B-catenin target (*Jag1*) and IFNGR genes in control and HFD MHC-II+ and MHC-II- ISCs (*n*=5). **F**. Organoid-initiating capacity of MHC-II+ and MHC-II- Lgr5-GFP^hi^ ISCs from HFD mice (*n*=5). **G**. Mean fluorescence intensity (MFI) of ISCs pulsed with either vehicle or DQ-Ovalbumin for 8 hours at 4C and 37C (*n*=5, 3 technical replicates per experiment). **H**. Weight of mice used for diet-induced obesity (left: Control and HFD, *n*=15) and leptin receptor deficiency models of obesity (right, Control and db/db, *n*=7) in the study. **I, J**. Frequency of MHC-II+ and MHC-II- ISCs in vehicle- and PPAR-delta agonist GW501516-treated mice by flow cytometry (**I**, *n*=3). Representative flow cytometry plots of MHC-II in vehicle- and PPAR-delta agonist GW501516-treated ISCs **(J)**. **K, L**. Frequency of MHC-II+ and MHC-II- ISCs in lean db/+ and obese db/db mice (**K**, *n*=3). Representative flow cytometry plots of MHC-II+ and MHC-II- ISCs (**I**, *n*=3). Unless otherwise indicated, data are mean ± s.d. from *n* independent experiments; \**P* □< □0.05, \*\**P* □< □0.01, \*\*\**P* □< □0.001 (Student’s *t*-tests).

**Supplementary Figure 2.**
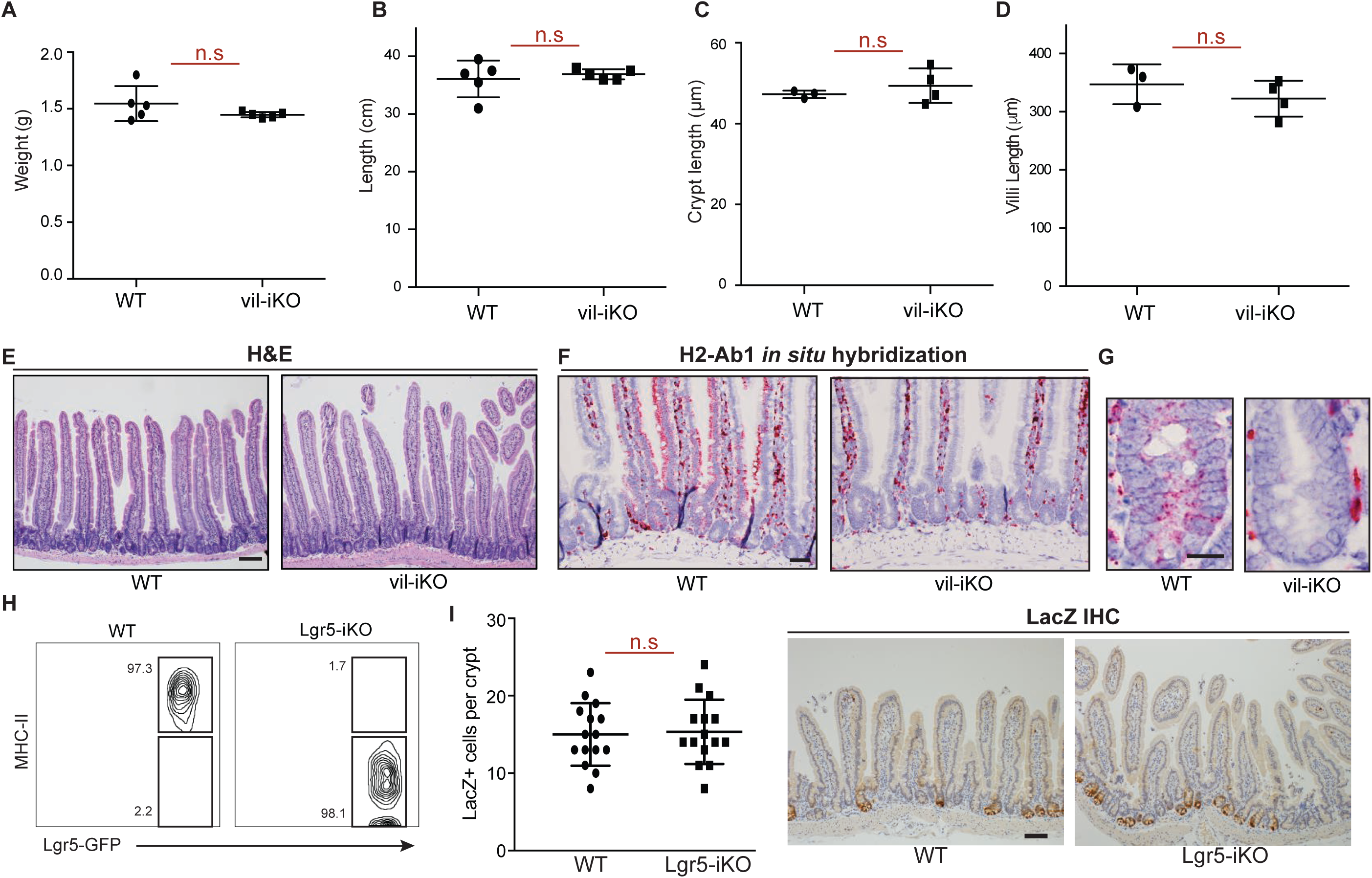
Intestine-specific deletion of MHC-II does not significantly alter intestinal physiology. **A**-**E**, Intestinal weight (**a**, *n*=5), length (**b**, *n*=5), crypt (**c**, *n*=5), and villi length (**d**, *n*=5) of MHC-II wild type (WT) and MHC-II^L/L^; Villin-CreERT2 (vil-iKO) mice one month after tamoxifen injection. Representative H&Es of WT and vil-iKO small intestine (**E**). **F, G**, *In situ* hybridization for *H2-Ab1* in the intestine (**F**) and representative images of intestinal crypts (**G**) in WT and vil-iKO mice (*n*=3). **H**. Frequencies of MHC-II+ and MHC-II- Lgr5-GFP^hi^ ISCs in WT and Lgr5-GFP MHC-II- deleted (Lgr5-iKO) by flow cytometry (*n*=5). **I**. Lineage tracing of LacZ+ cells in the intestinal crypt after deletion of MHC-II in ISCs (left). Representative images of LacZ immunostain in WT and Lgr5-iKO small intestine (right). Unless otherwise indicated, data are mean □± □s.d. from *n* independent experiments; \**P* □< □0.05, \*\**P* □< □0.01, \*\*\**P* □< □0.001 (Student’s *t*-tests). Scale bars, 100 □μm (**E**), 50 □μm (**F, I, K**) and 20 □μm (**G, I**, (insets)).

**Supplementary Figure 3.**
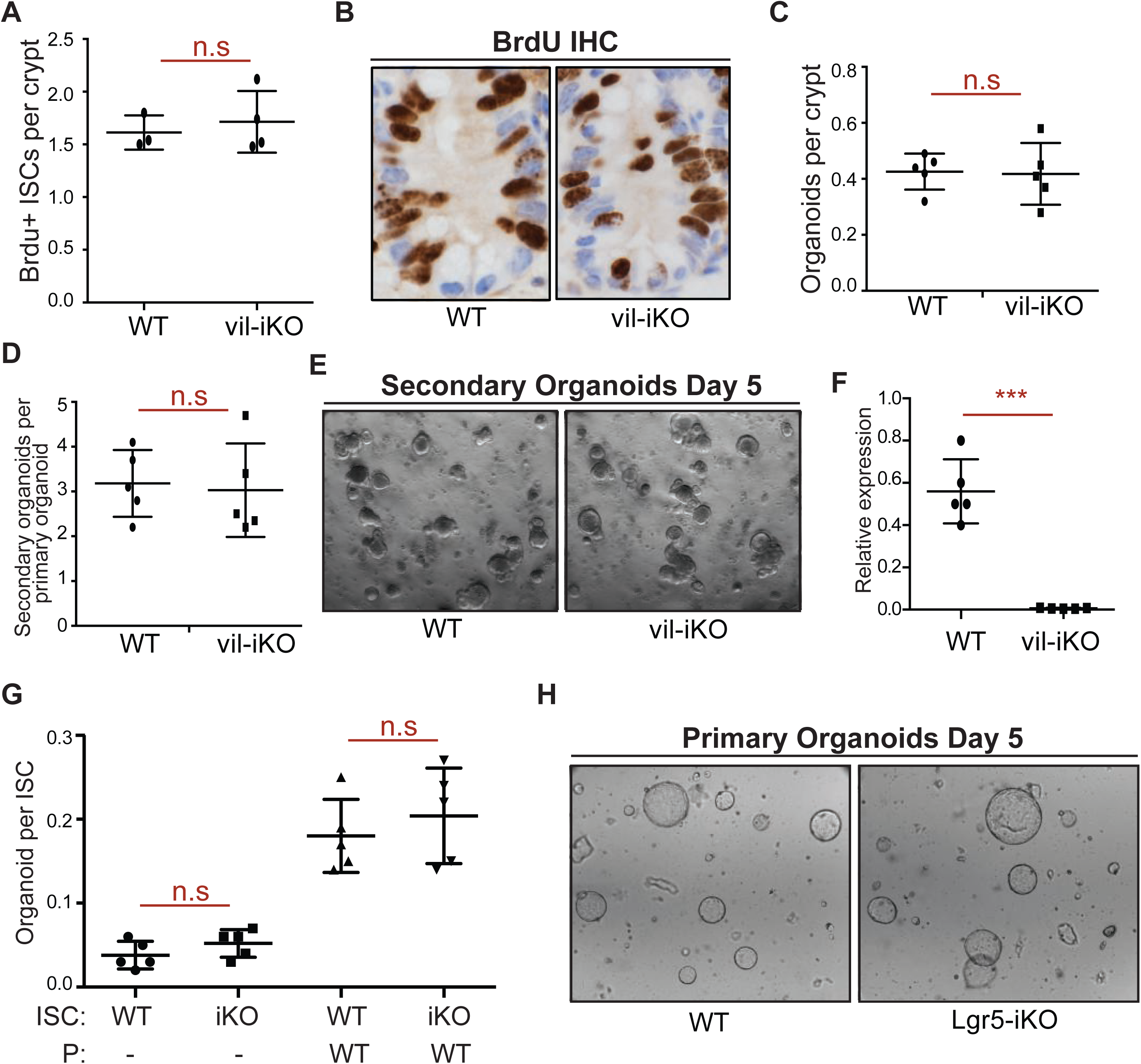
Intestine-specific deletion of MHC-II does not alter the organoid forming capacity of ISCs. **A, B**. Number of bromodeoxyuridine (BrdU)+ crypt base columnar cells after a 4-hour pulse in MHC-II^L/L^; Villin-CreERT2 (vil-iKO) mice after tamoxifen administration (WT: *n*=3, vil-iKO: *n*=4). Representative images of BrdU immunostain in proximal small intestinal crypts (**B**). **C**-**E** Organoid-initiating capacity of WT and vil-iKO crypts (**C**, *n=*5). Number of secondary organoids per dissociated crypt-derived primary organoid (**D**, *n=*5). Representative images of day-5 WT and vil-iKO primary organoids (**E**). **F**. Relative expression of MHC-II in dissociated WT and vil-iKO primary organoids at day 5 (*n=*5). **G, H**. Organoid-initiating capacity of ISCs from WT and MHC-II^L/L^; Lgr5-EGFP-IRES-CreERT2 (Lgr5-iKO) mice with and without Paneth cells (P) from WT mice (*n=*5). Representative images of organoids derived from WT and Lgr5-iKO ISCs co-cultured with WT Paneth cells five days after seeding (**H**). Unless otherwise indicated, data are mean □± □s.d. from *n* independent experiments; \**P* □< □0.05, \*\**P* □< □0.01, \*\*\**P* □< □0.001 (Student’s *t*-tests). Scale bars, 100 μm (**E, H**) and 20 μm (**B**).

**Supplementary Figure 4.**
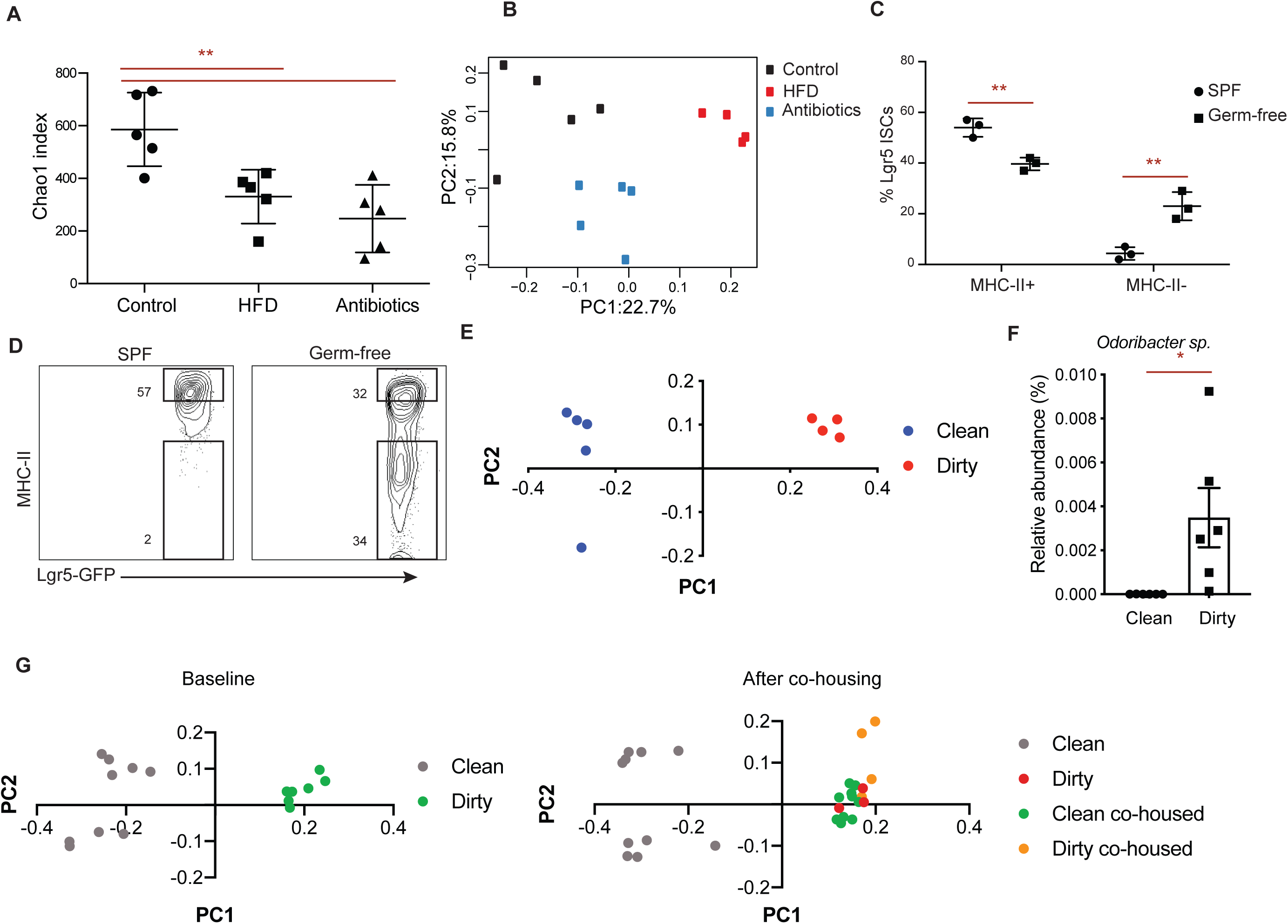
Microbial regulation of MHC-II expression in ISCs. **A**. Chao1 index of microbial diversity in Control, HFD, and mice treated with antibiotics for 3 months (*n*=5). **B**. Principle component analysis of microbial diversity in Control, HFD, and antibiotic-treated mice (*n*=5). **C, D**. Frequencies of MHC-II+ and MHC-II- ISCs in specific-pathogen free (SPF) and germ-free mice by flow cytometry (*n*=3). Representative flow cytometry plots of MHC-II in SPF and germ-free ISCs **(D)**. **E**. Principle component analysis of microbial diversity in mice housed in clean room or dirty room (*n*=6). **F**. Relative abundance of *Odoribacter sp*. in mice housed in clean room or dirty room (*n*=6). **G**. Principle component analysis of microbial diversity at baseline level before co-housing experiment (left) and 10 days after co-housing clean mice with dirty mice in the dirty room. Unless otherwise indicated, data are mean □± □s.d. from *n* independent experiments; \**P* □< □0.05, \*\**P* □< □0.01, \*\*\**P* □< □0.001 (Student’s *t*-tests).

**Supplementary Figure 5.**
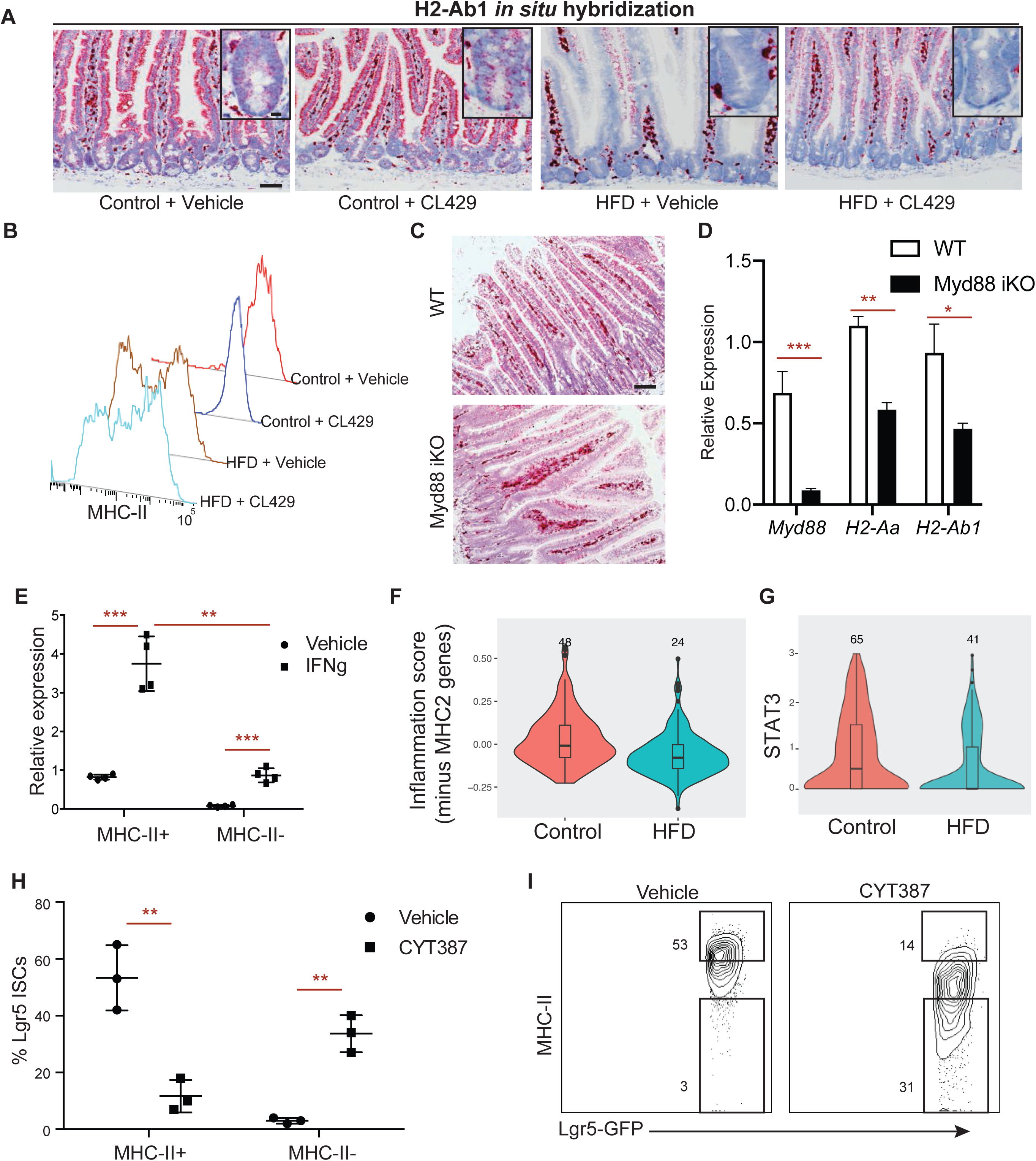
Regulation of MHC-II expression in ISCs by PRR and Jak/Stat signaling. **A, B**. *In situ* hybridization for H2-Ab1 in control and HFD mice treated with vehicle- and CL429-treated mice in proximal small intestine (**A**, *n=*3). Representative histogram plots of MHC-II in ISCs in control and HFD mice treated with vehicle and CL429 (**B**, *n*=3). **C**. *In situ* hybridization for H2-Ab1 in WT and Myd88 KO mice (*n*=4). **D** Relative expression of *Myd88* and MHC-II genes (*H2-Aa* and *H2-Ab1*) in the intestine from WT and Myd88 KO mice (*n=*5). **E** Relative expression of MHC-II (*H2-Ab1*) in MHC-II+ and MHC-II- ISCs isolated from HFD mice with or without IFNg stimulation (*n*=4). **F** Violin plots demonstrating the levels of IFNg-induced genes excluding MHC-II pathway genes in control and HFD ISCs by scRNA-seq. **G** Violin plots demonstrating the levels of STAT3 in control and HFD ISCs by scRNA-seq. **H, I**. Frequencies of MHC-II+ and MHC-II- ISCs in vehicle- and Jak/Stat inhibitor CYT387-treated mice (*n*=3). Representative flow cytometry plots of MHC-II in vehicle- and CYT387-treated ISCs (**I**). Unless otherwise indicated, data are mean □ ± □ s.d. from *n* independent experiments; \**P* □ < □ 0.05, \*\**P* □ < □ 0.01, \*\*\**P* □ < □ 0.001 (Student’s *t*-tests). Scale bars, 50 μm (**A, C**) and 20 μm (insets, **A**).

**Supplementary Figure 6.**
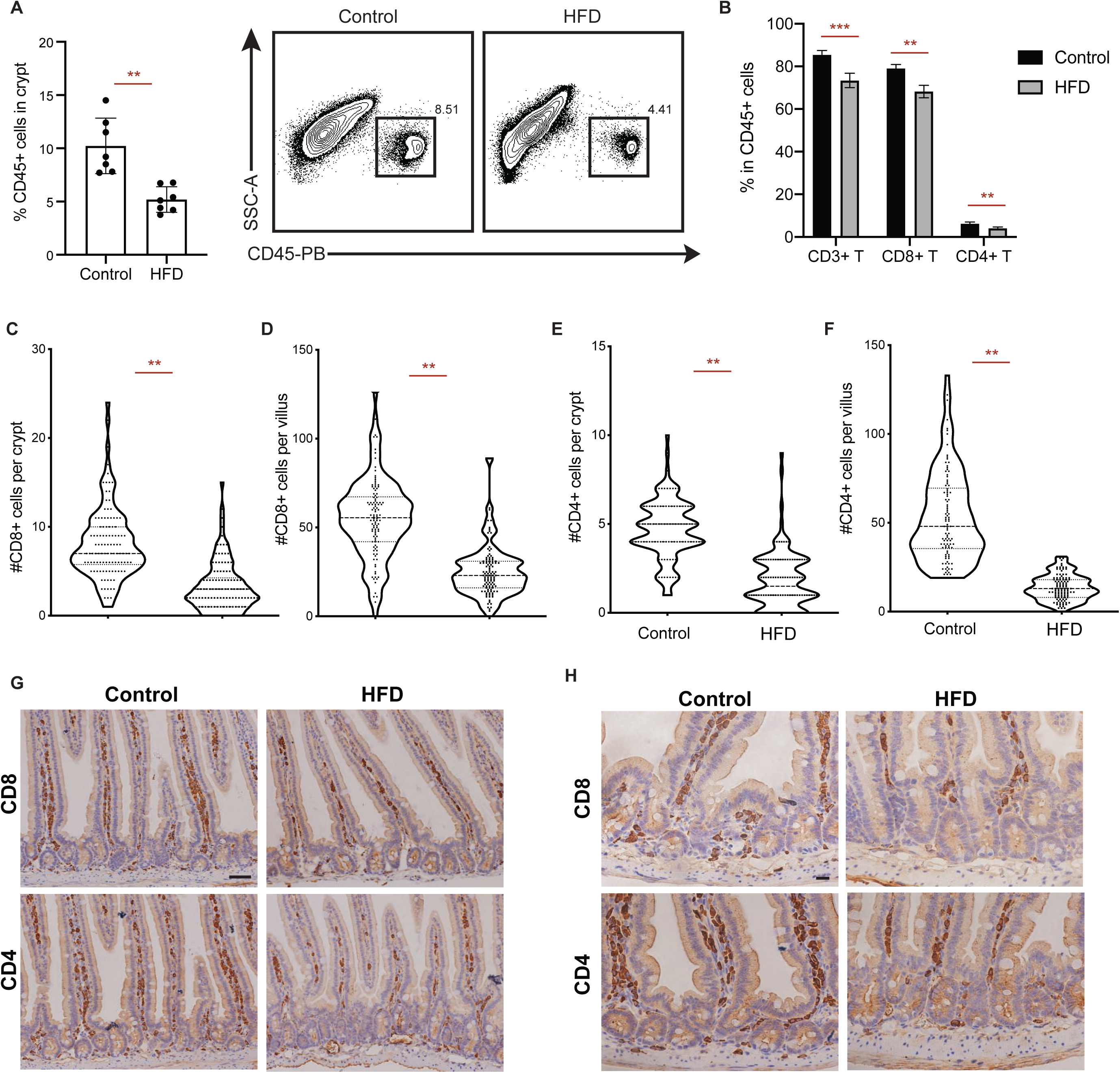
The effect of HFD on intestinal immune cells. **A**. Frequency of CD45+ cells in intestinal crypts isolated from control or HFD mice (n=7). **B**. Frequency of CD3+ T cells, CD8+ T cells and CD4+ T cells among CD45+ cells in intestinal crypts isolated from control or HFD mice (*n*=5). **C-H**. Numbers of CD8+ cells (**C**, crypt; **D**, villus) and CD4+ cells (**E**, crypt; **F**, villus) in the intestines of control or HFD mice. Representative images of CD4 and CD8 immunostaining in control or HFD intestines (**G, H**).

**Supplementary Figure 7.**
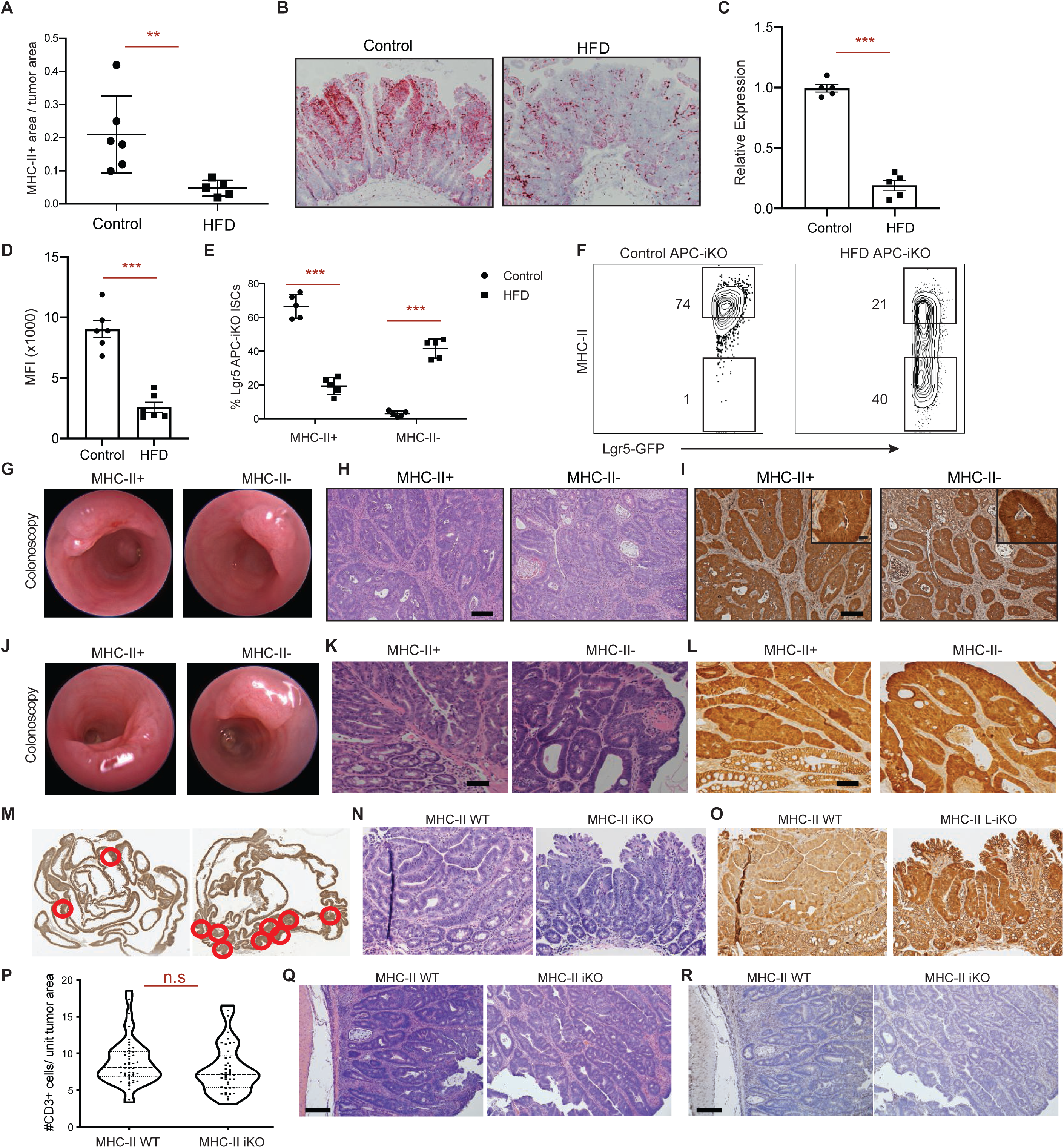
The effect of ISC MHC-II expression on tumor formation. **A, B**. Fraction of *H2-Ab1*+ tumor areas per total tumor area by single-molecule *in situ* hybridization in control and HFD Lgr5+ *Apc*-null mice 14 days post tamoxifen injection (Control, *n*=6; HFD, *n*=5). Representative images of H2-Ab1 *in situ* hybridization in control and HFD tumors from Lgr5+ *Apc*-null mice (**B**). **C**. Relative expression of MHC-II (*H2-Ab1*) in Lgr5+ *Apc*-null pre-malignant ISCs isolated from control or HFD mice 3 days post tamoxifen injection (*n*=5). **D**. Mean fluorescence intensity (MFI) of MHC-II in Lgr5+ *Apc*-null pre-malignant ISCs isolated from control or HFD mice 3 days post tamoxifen injection (*n*=6). **E, F**. Frequency of MHC-II+ and MHC-II- Lgr5-GFP^hi^ *Apc*-null ISCs in control and HFD mice by flow cytometry (**E**, *n*=5). Representative flow cytometry plots of MHC-II in HFD *Apc*-null ISCs (**F**). **G**-**I**. Characterization of orthotopically transplanted MHC-II+ and MHC-II- *Apc*-null ISCs-derived tumors three months after transplantation into immunocompetent syngeneic hosts. Optical colonoscopy images (**G**), Hematoxylin and eosin (**H**) and beta-catenin immunostain (**I**) of tumors. **J**-**L**. Characterization of orthotopically transplanted MHC-II+ and MHC-II- *Apc*-null ISCs-derived tumors three months after transplantation into immunodeficient Rag2-KO hosts. Optical colonoscopy images (**J**), H&E (**K**) and beta-catenin immunostain (**L**) of tumors. **M-O**. Tumors in Lgr5-Cre+ APC^+/-^ MHC^+/-^ (MHC-II WT) and Lgr5-Cre+ APC^+/-^ MHC^-/-^ (MHC-II iKO) mice. Representative images of tumors from Lgr5-Cre+ APC^+/-^ MHC^+/-^ (**M**, left) and Lgr5-Cre+ APC^+/-^ MHC^-/-^ (**M**, right) mice. Representative H&E (**N**) and beta-catenin immunostain (**O**) images from tumors in Lgr5-GFP+ APC^+/-^ MHC^+/-^ and Lgr5-GFP+ APC^+/-^ MHC^-/-^ mice. **P-R**. Numbers of CD3+ T cells per tumor area (**P**). Representative H&E (**N**) and CD3 immunostain (**O**) images from tumors.

## Notes

### Competing Interest Statement

The authors have declared no competing interest.

